# Dynamic Insights into Hsp90 Inhibitor Binding: Uncovering Intermediate States and Conformational Plasticity

**DOI:** 10.1101/2024.10.19.619188

**Authors:** Mohammad Sahil, Jaya Krishna Koneru, Jagannath Mondal

**Affiliations:** Tata Institute of Fundamental Research Hyderabad, 36/P Gopanapalli Village, TS-500046, India

## Abstract

Heat shock protein 90 (Hsp90) is a vital molecular chaperone involved in protein folding, stabilization, and activation, making it a key target in cancer therapy. Targeting the N-terminal domain of Hsp90 (N-Hsp90) has emerged as a potential strategy for developing anticancer therapies due to its involvement in the folding and function of oncoproteins. However, the precise molecular details of how inhibitors bind to N-Hsp90 remain elusive due to inherent conformational plasticity involving *loop-in* and *loop-out* states of loop-4 (L4). In this work, we utilized unbiased molecular dynamics (MD) simulations coupled with Markov state modeling (MSM) and machine learning techniques to investigate the binding dynamics of the inhibitor Geldanamycin (GDM). Our findings reveal a complex two-stage binding mechanism involving the formation of a non-native intermediate state prior to the final bound state. We demonstrate that GDM binding predominantly stabilizes the *loop-out* conformation of N-Hsp90, challenging prior beliefs and suggesting that *both* conformational selection and induced fit mechanisms are involved. Through a detailed residue-level analysis using Random Forest based ML classifiers, we quantify the relative contributions of these mechanisms, highlighting the dynamic interplay between Hsp90’s conformational states during inhibitor binding. These findings enhance our understanding of Hsp90’s conformational plasticity and could inform the design of more effective Hsp90 inhibitors.

## Introduction

Heat shock protein 90 (Hsp90) is one of the most abundant cytosolic proteins in eukaryotes, accounting for 4-6% of the total cytosolic protein content under stress conditions.^1^ As a ubiquitous molecular chaperone, Hsp90 plays a crucial role in facilitating the folding and stabilizing the higher-order assemblies of a wide range of client proteins. ^2,3^ These clients rely on Hsp90 to maintain their functional conformations, especially under stress conditions.^4^ Structurally, Hsp90 consists of three functional domains: the N-terminal domain (N-Hsp90), the middle domain, and the C-terminal domain.^5^ Hsp90 functions as a dimer, with the C-terminal domains forming a V-shaped dimeric structure that positions the client proteins between the two middle domains. ^3^ The activation of Hsp90 is tightly regulated by conformational changes in the N-terminal domain, which houses an ATP-binding site. ATP binding induces conformational changes in both the N-terminal and middle domains, thereby promoting ATP hydrolysis and activating the chaperoning function of Hsp90. ^6^

Hsp90’s chaperoning activity is essential not only for maintaining cellular homeostasis under normal and stressed conditions but also for supporting the survival of cancer cells, where elevated levels of Hsp90 are often observed.^7^ The uncontrolled growth and high stress levels in cancer cells necessitate the chaperoning function of Hsp90 to assist in the folding and stabilization of oncoproteins, while simultaneously inhibiting stress-induced apoptosis.^1^ The critical role of Hsp90 in tumor cell survival has been established across various cancer types, including both solid tumors and hematological malignancies. Inhibiting or knocking out Hsp90 has been shown to suppress tumor growth, making N-Hsp90 a promising target for anti-cancer therapies.^8^

Given its potential in cancer treatment, numerous Hsp90 inhibitors have been developed and tested.^1^ Geldanamycin (GDM), a natural compound extracted from *Streptomyces hygroscopius*, was one of the first inhibitors to enter clinical trials. ^9^ Although GDM effectively inhibited Hsp90, its clinical development was halted due to unacceptable toxicological effects observed in vivo. However, several derivatives of GDM, such as 17-AAG and 17-DMAG, have since advanced to clinical trials. Another notable class of Hsp90 inhibitors includes Radicicol and its derivatives.^1^ In addition to natural inhibitors, numerous synthetic inhibitors representing various chemical classes such as purines, pyrazoles, triazines, quinolines, coumarins, isoxazoles, and hybrids-are under investigation.^10^ To date, more than 100 inhibitors from these classes have been extracted or synthesized, but none have yet been finalized for clinical use.^11^ Despite this, ongoing research continues to focus on the development of new derivatives, particularly for GDM, with over 40 derivatives currently known and more in development.^12–15^

Despite the significant interest in Hsp90 inhibitors, the molecular and atomistic basis of their binding mechanisms remains incompletely understood. Hundreds of crystal structures of N-Hsp90 bound to ATP or various inhibitors have been determined, consistently revealing a single binding site for substrates or inhibitors (figure 1A). While crystallography has provided valuable insights into the bound state, NMR studies of GDM suggest a two-step binding mechanism involving an intermediate state via an induced fit mechanism. ^16^ Conversely, a single-step binding mechanism has been proposed for the GDM derivative 17-DMAG,^17^ potentially indicating a conformational selection route for inhibitor binding. As discussed below, the substrate-binding mechanism is further complicated by the strong conformational heterogeneity present in the Hsp90 binding pocket.

**Figure 1:**
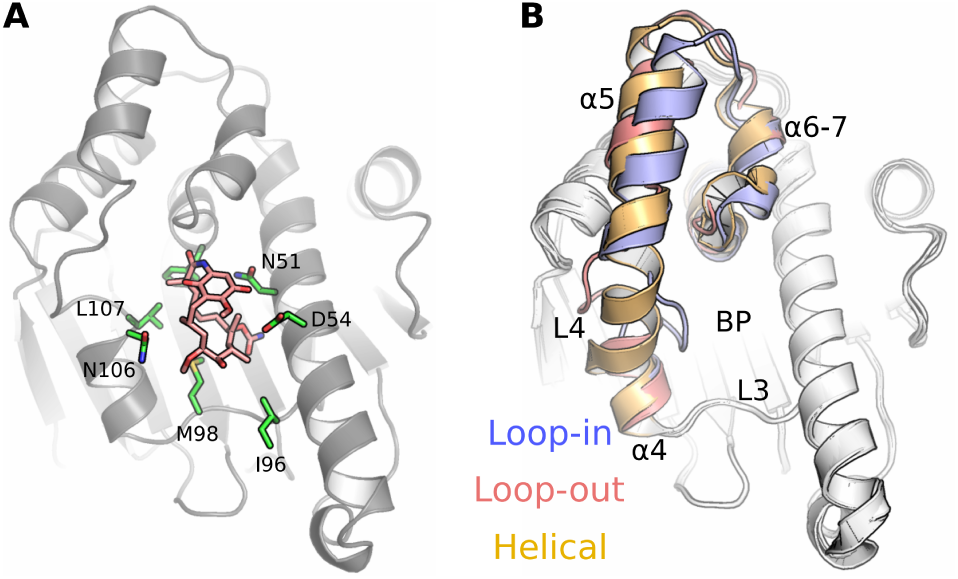
Conformational plasticity in Hsp90. (A) Native bound state of GDM as observed for crystallography (pdb id: 1YET). (B) Three states of N-Hsp90 highlighted for residues 100-144, corresponding to pdb ids 1YER (loop-in), 1YES (loop-out) and 5J20 (helical). BP represents binding pocket.

A key aspect of N-Hsp90 is the high conformational plasticity observed at loop-4 (L4 or the lid), which can broadly be classified into three states: *loop-in* (a closed conformation, e.g., PDB ID: 1YER), *loop-out* (an open conformation, e.g., PDB ID: 1YES), and an intermediate *helical* conformation (e.g., PDB ID: 5J20, figure 1).^18^ Prior NMR^19^ and MD^20–22^ studies have provided further evidence for the existence of multiple conformations of Hsp90 in its free form, prior to any binding event. Conformational plasticity is known to affect biomolecular recognition,^23^ and it is crucial for inhibitor binding in Hsp90, with the entropic contribution of loop-4 playing a significant role.^18^ Most Hsp90 inhibitors exhibit specificity for particular protein conformations, binding preferentially to one of the L4 states, and are thus referred to as either ‘loop binders’ or ‘helical binders.’ The substrate of interest in this study, the parent molecule GDM, specifically binds to the loop-out conformation, as observed in substrate-bound crystal structures (PDB ID: 1YET).^24^ Despite extensive studies focusing on the end states of Hsp90 (i.e., apo or bound states),^25,26^ a detailed understanding of the binding process and its interplay with the conformational heterogeneity of Hsp90 is still lacking. For instance, whether GDM selects a pre-existing loop-out state (conformational selection) or induces the loop-out state, or a combination of both, remains unclear.

In this work, we aim to elucidate the role of conformational heterogeneity in the binding of GDM to Hsp90 by simulating the recognition process of GDM within its native cavity. To this end, we conduct extensive unbiased molecular dynamics (MD) simulations to track the search for the native binding pose of GDM in Hsp90, complemented by adaptively sampled trajectories. We employ a comprehensive Markov state modeling (MSM) approach to characterize the dynamic binding process of GDM to Hsp90, identifying a two-stage recognition mechanism facilitated by a binding intermediate. Additionally, a follow-up MSM analysis of both the apo and native substrate-bound states sheds light on the conformational heterogeneity near the L4 region, revisiting the debate on conformational selection versus induced fit. Finally, we resolve this debate by applying a Random Forest classifier to the simulated trajectories, quantifying the contributions of both mechanisms at play in the binding process.

## Methods

### Molecular Dynamics Simulations

Four sets of all-atom, unbiased molecular dynamics (MD) simulations were conducted for the N-terminal domain of human Hsp90 (N-Hsp90, residue 9-236). A total of 38 replicates, accumulating to 62 *µ*s of simulation time, were performed (Table 1).

**i) Unbiased Binding Kinetics Simulations:** In the first set, unbiased binding kinetics simulations were performed to investigate the binding of Geldanamycin (GDM) to the apo form of N-Hsp90. GDM molecules, initially placed in the solvent phase, were allowed to explore the environment around the apo N-Hsp90 in search of the native binding pocket. The starting configuration of human Hsp90*α* was taken from previously solved crystal structures of the apo form (PDB ID: 1YES, loop-out conformation, figure 2A). The all-atom MD system comprised N-Hsp90 with residues protonated at neutral pH, neutralized with 180 mM KCl, and solvated in a TIP3P water box with a 10 *AÅ* buffer around the protein. The CHARMM36^27^ force field was used to parameterize the protein. For the binding simulations, four GDM molecules, corresponding to a concentration of 10 mM, were randomly placed in the vicinity of the apo protein. The force field parameters for GDM were derived using GAAMP.^28^ All simulations were preceded by 250 ps of NVT and NPT equilibrations, em-ploying the v-rescale thermostat^29^ (310.15 K) and the Parrinello-Rahman^30^ barostat. The simulations were executed using GROMACS 20xx,^31^ with non-bonded interactions calculated via Particle Mesh Ewald (PME)^32^ summation and the Verlet^33^ scheme within a 1.2 nm cutoff. Hydrogen bonds were constrained using the LINCS^34^ and SETTLE^35^ algorithms.
**ii) Adaptive Sampling for Markov State Model Development:** To comprehensively sample the ligand-binding phase space for subsequent Markov state model (MSM) development (described later), multiple configurations corresponding to intermediate states were selected from the long binding trajectories. Shorter simulations (100 ns each) were then adaptively initiated from these intermediate states.
**iii) and iv) Comparative Simulations of Apo and Bound Forms:** For a comparative assessment of the conformational heterogeneity of Hsp90 in its apo and native-bound forms, additional simulations were performed starting from crystal structures corresponding to the apo form (PDB IDs: 1YES, 1YER) and the bound form (PDB ID: 1YET).

**Figure 2:**
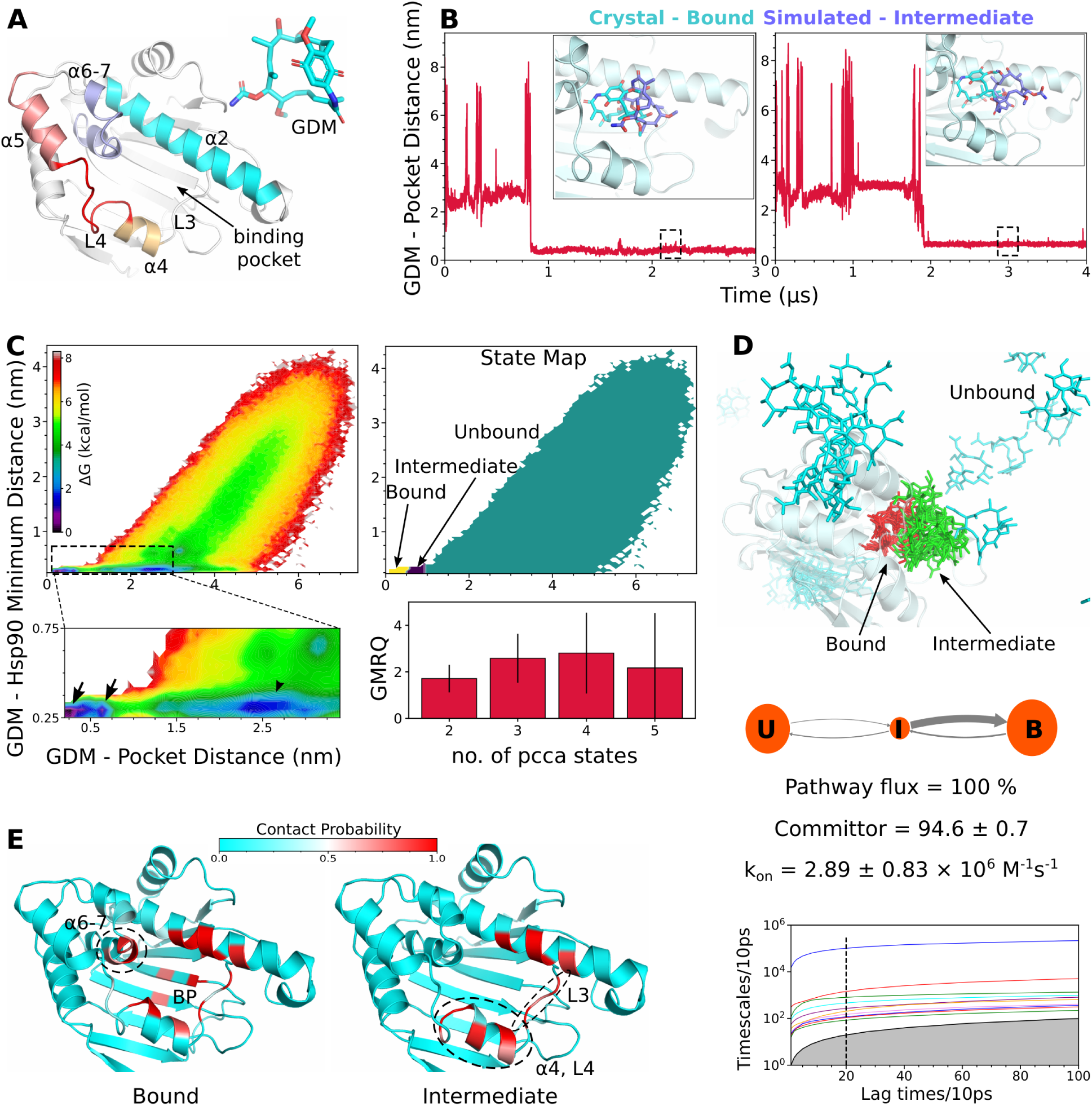
Three state inhibitor binding to N-Hsp90. (A) The N-terminal domain (residue 9-236) of human-Hsp90 and parent inhibitor geldanamycin (GDM, unbound state) structure. (B) Ligand distance profile in two binding simulations, leading to the formation of putative intermediate state. Snapshot represent the conformation from dashed inset. (C) Free energy surface (FES) of binding process, corresponding state map achieved by three state pcca classification and GMRQ scores for MSM coarse grained at different number of states. Arrow and arrowhead in FES plot indicates minimas. (D) The three state MSM of GDM binding to N-Hsp90 involving unbound (U), intermediate (I) and bound (B) states. Below: Binding observables estimated by three state MSM and implied timescales indicating three states at lag time 20 steps. (E) The GDM-protein contact probability map corresponding to intermediate and bound states.

**Table 1:** Simulation replicates and time scales.

| **simulation** | **number of replicates** | **total simulation time (μs)** |
| --- | --- | --- |
| Unbiased binding simulations | 8 | 20.97 |
| Adaptive sampling | 20 | 2 |
| Apo state | 6 | 23.26 |
| Bound state | 4 | 16 |

### Development of Markov State Models

Three independent Markov state models (MSMs)^36^ were constructed using different sets of unbiased MD data to investigate: (i) the GDM binding process to its native pocket in Hsp90, and (ii) and (iii) the loop dynamics near the L4 region of Hsp90 in both apo and bound states.

For the MSM development focused on the binding process, two collective variables (CVs) were selected: the center-of-geometry distance between GDM and the binding pocket (residues 51, 54, 58, 98, 102, 106, 107, 135, 138, figure S1) and the minimum distance be-tween GDM and the protein. These CVs were used to discretize the simulated trajectories as the initial step in constructing the MSM. Notably, an attempt to use the GDM-protein contact matrix as a clustering metric did not successfully resolve the states.

For the MSM characterizing the loop dynamics involving the L4 region (residues 106-115), minimum distances between each loop residue and every third residue of the rest of the protein (resulting in a 630-dimensional CV array) were considered. To reduce the dimensionality of the input CVs for the loop dynamics MSM, time-structured independent component analysis (TICA) was employed to extract the two slowest motions. These were subsequently clustered into microstates using the k-means algorithm.

Table 2 lists the relevant hyperparameters used in the respective MSMs. The final MSMs were built over a range of lag times and were coarse-grained into 2-7 states using the PCCA (Perron Cluster Cluster Analysis) algorithm (ITS plots in figures 2 and S4). The final settings were chosen based on the convergence of PCCA states across all lag times (Table 2). Pathways and associated transition timescales were calculated using transition path theory.

**Table 2:** Hyperparameters used in markov state models.

| MSM | data size (μs) | tica | clusters | lag time (ns) | pcca states |
| --- | --- | --- | --- | --- | --- |
| binding process | 38.4 | - | 300 | 0.2 | 3 |
| loop dynamics - Apo | 23.26 | 5 ns | 50 | 0.2 | 6 |
| loop dynamics - Bound | 16 | 5 ns | 50 | 0.2 | 4 |

The MSMs were constructed and analyzed using the Python libraries `pyemma`^37^ and `msmbuilder`,^38^ with GMRQ scores employed for model evaluation.

### Ligand Interactions

Ligand interactions with Hsp90 in both intermediate and bound states were quantified by calculating the probability of contact between specific residues and the GDM ligand. Contacts were defined as a minimum distance *≤* 0.51 nm between a residue and the ligand. These probabilities were computed over all frames from the intermediate (binding simulations) and substrate-bound trajectories.

### Perturbation Response Scanning

The propensity for interconversion between the loop-in and loop-out states was assessed using perturbation response scanning (PRS)^39^ based on linear response theory, as implemented in the `md-task`^40^ utility. The crystal structures of the loop-in (PDB ID: 1YER) and loop-out (PDB ID: 1YES) conformations were used as reference end states. All simulation trajectories of the apo form of Hsp90 (Table 1) were employed to construct the variance-covariance matrix. A total of 1000 perturbation forces were applied to each residue. PRS was performed for two transitions: ‘loop-in to loop-out’ and ‘loop-out to loop-in.’

### Pocket Volume Estimation

The 3D pocket volume of the N-Hsp90 active site was estimated using structures aligned from the apo and bound state simulation ensembles. The `MDpocket`^41^ utility was employed to sample the pocket architecture, utilizing the Voronoi tessellation algorithm.^42^ The pocket volume was calculated as the cumulative volume of alpha spheres detected at a specific site in more than 50% of the simulation ensemble. In approximately 4-5% of frames, the pocket could not be detected due to side-chain flexibility or limitations of `MDpocket`; such frames were excluded from analysis. The pocket volumes of the detected frames were bootstrapped for 1000 iterations to smooth the probability density curve (with the mean unchanged). The apo state pocket volumes were further classified into loop-out, loop-in, and helical states of loop-4, according to MSM-derived macrostates of the apo state, corresponding to [LO], [LI, LI1, LI2, LI3], and [H] macrostates, respectively.

### Random Forest Training

Two sets of input features were calculated from all simulations of apo and bound states (Table 1): (i) native contacts, defined as minimum distances between all residue pairs with *C_α_ − C_α_* distances within 10 *Å* in the starting structure, and (ii) loop-4 distances, defined as minimum distances between each loop-4 residue (106-115) and every third residue of the rest of the protein (also used in MSM). The two feature sets contained 1194 and 630 dimensions, respectively. Two random forest (RF) classifiers were trained on each feature set.

For the first classifier, features from only apo simulation trajectories were labeled according to six MSM states, which were further coarse-grained into three states: loop-out (0), loop-in (1), and helical (2). The second classifier used features from both apo and bound simulations, labeled as apo (0) and bound (1). Each RF model consisted of 1000 estimators, using Gini impurity as a measure of leaf purity. Fifty independent cross-validated RF models were trained by randomly selecting 70% of the training data, with the remaining 30% used for validation based on accuracy (Eq. 1) and F1 scores (Eq. 2). The final output of successfully trained classifiers was the feature importance scores (*F_imp_*, Eqs. 3-4).

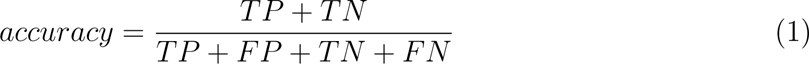

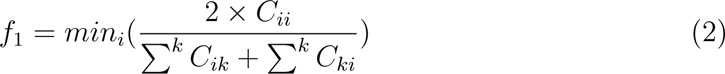

where, TP, FP, TN, and FN are true positive, false positive, true negative and false negative components of a k (k=i) dimensional confusion matrix estimated on test data.

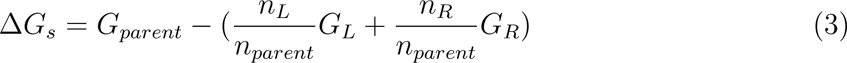

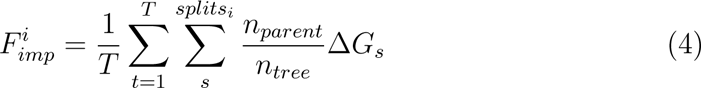

given that an arbitrary parent node with *G_parent_* impurity and *n_parent_* datapoints is split into child nodes having *G_R_* and *G_L_* impurities and *n_R_* and *n_L_* datapoints. *F_imp_* is weighted mean (by datapoints in tree (*n_tree_*) and *n_parent_*) of information gain (Δ*G_s_*) over all splits in all estimators (*T*).

## Results and Discussion

### Inhibitor binding to N-Hsp90 occurs via a potential intermediate state

We began by investigating the binding mechanism of the inhibitor Geldanamycin (GDM) to N-Hsp90 through unbiased molecular dynamics (MD) simulations. Unbiased binding simulations^23,43,44^ are widely recognized for their ability to recover binding mechanisms with atomistic detail, allowing ligand molecules in the unbound state to search for binding modes and ultimately bind to the native pocket as observed in crystal structures. In this study, the simulation box was filled with a single copy of N-Hsp90 (residues 9-236, figure 2A), randomly surrounded by four GDM molecules (corresponding to a 10 mM concentration), positioned away from the Hsp90 target in the unbound state. Eight independent replicates were performed, each ranging from 0.5 to 6 *µ*s in duration. The center of geometry (COG) distance between GDM and the binding pocket was used as a metric to monitor the binding process.

Despite our efforts, the GDM molecules did not attain the crystal-bound pose (PDB ID: 1YET) in any of the binding simulations, which is not entirely surprising given the slow binding nature of the large GDM inhibitor-a timescale potentially beyond the reach of conventional MD simulations. However, in two of the eight binding simulations, a stable binding mode near the native pocket was observed (figure 2B). We confirmed that this pose was not transient or unstable, as it persisted over multiple microseconds of simulation. A closer examination suggests that, due to its proximity to the binding pocket, this observed binding mode may serve as a potential intermediate in the binding mechanism. This observation aligns with previous biochemical assays that indicated a two-step binding mechanism^16^ involving an intermediate state.

To further elucidate the binding mechanism and confirm the existence of this intermediate state as a long-lived metastable state, we constructed a Markov state model (MSM). In addition to the two binding trajectories shown in figure 2B, we performed four bound simulations to sample the crystal-bound pose. The unsuccessful binding trajectories were also included to sample the unbound region. To further explore the transitions of the poses observed in figure 2B, we carried out 20 adaptive sampling simulations, each lasting 100 ns, spawned from the parent binding trajectories. The MSM was built using two collective variables: the COG distance between GDM and the binding pocket, and the minimum distance between GDM and Hsp90.

The resulting two-dimensional free energy surface (FES), reweighted by MSM-derived equilibrium populations of microstates, reveals two proximal, deep minima, which likely correspond to the native-bound and metastable intermediate states (figure 2C). Additionally, a less deep minimum is observed at approximately 2.5 nm, suggesting the presence of an additional non-native bound state.

To verify the convergence of the key macrostates within the ensemble, we first divided the free energy surface (FES) into two metastable states using the PCCA algorithm applied to Markov state models (MSMs) constructed at various lag times (10, 20, … 60 timesteps) (figure S2). The resulting state maps were consistent across all lag times, indicating stability. These two states corresponded to the bound (both crystal-bound and simulation-observed binding mode) and unbound (remaining states) conformations.

Next, we explored the MSM with three PCCA states, which also produced converged state maps at all lag times (figure S2). In this MSM, the three states were identified as bound (crystal-bound), potential intermediate (simulation binding mode), and unbound (remaining states). To further investigate the additional minimum observed at 2.5 nm, we constructed MSMs with four and five PCCA states. However, these models did not yield converged state maps (figure S2). Although the GMRQ score of the three-state MSM was higher than those of other models, except for the four-state MSM, the fourth state was not physically meaningful in the context of binding (figure S2). Therefore, we concluded that GDM binding to Hsp90 involves three primary states: unbound, bound, and intermediate (figures 2D, S3, S4).

To estimate the possible binding pathways, we examined the transition from unbound to bound states. The analysis revealed a single dominant pathway, corresponding to the sequence unbound *→* intermediate *→* bound, with 100% flux (figure 2D). This finding suggests that GDM binding to Hsp90 proceeds via a single non-native intermediate, with direct binding from unbound to bound being infeasible. The committor probability of the intermediate state was calculated to be 0.946*±*0.007, indicating that once GDM transitions from the unbound to the intermediate state, there is a 94% probability of proceeding to the bound state. Overall, these simulations suggest that GDM binding to Hsp90 occurs through a potential intermediate, in agreement with NMR-based observations.^16^

Although the crystal-bound state and the simulation-observed intermediate exhibit minor positional differences, the two states are significantly different in terms of residue interactions (figure 2E). A comparison of contact probabilities revealed that the bound state is characterized by specific ligand-receptor contacts buried within the basal beta sheets of the binding pocket (BP) and helices *α*6-*α*7, while the intermediate state shows increased surface contacts, particularly with loop-3 (L3) and helix *α*4.

### Substrate Recognition Depicts a Complex Picture of the Underlying Conformational Space of N-Hsp90

After establishing the two-stage substrate binding mechanism in Hsp90, we next focused on characterizing the key changes in the protein’s conformational space upon substrate binding to the native pocket, relative to the apo form. Adjacent to the native binding pocket of Hsp90, the loop-4 (L4, between *α*4 and *α*5) region is known to exhibit a remarkably high degree of conformational heterogeneity.^18,19^ The L4 region, encompassing residues 106-115, has been broadly categorized into three major conformations: loop-in (L4 facing inside/towards the binding pocket), loop-out (L4 facing away from the binding pocket), and helical (an intermediate state where L4 adopts a helical conformation, merging *α*4 and *α*5 into one helix, figure 1). Notably, the substrate-bound crystal structure of GDM depicts the *loop-out* conformation of L4.

While crystallographic poses often represent a single conformation for each inhibitor, proteins are now increasingly recognized as ensembles of structures.^45^ A recent study revealed the complex behavior of conformational heterogeneity in the L4 region of N-Hsp90.^18^ As the next logical step in understanding the GDM binding process in N-Hsp90, we aimed to elucidate the conformational changes in the L4 region induced by GDM binding. To this end, independent simulated ensembles were generated for the ‘apo’ (N-Hsp90 alone) and ‘bound’ (GDM-bound N-Hsp90) states, and two independent MSMs were constructed for each ensemble. To capture all possible conformations of the L4 region, a large array of distances between L4 residues (106-115) and every third residue of the rest of N-Hsp90 was considered, which was subsequently dimensionally reduced via TICA for MSM construction (Methods). For the apo simulations, the MSM converged to six PCCA states, whereas for the bound simulations, four PCCA states were identified (figures 3, S4).

**Figure 3:**
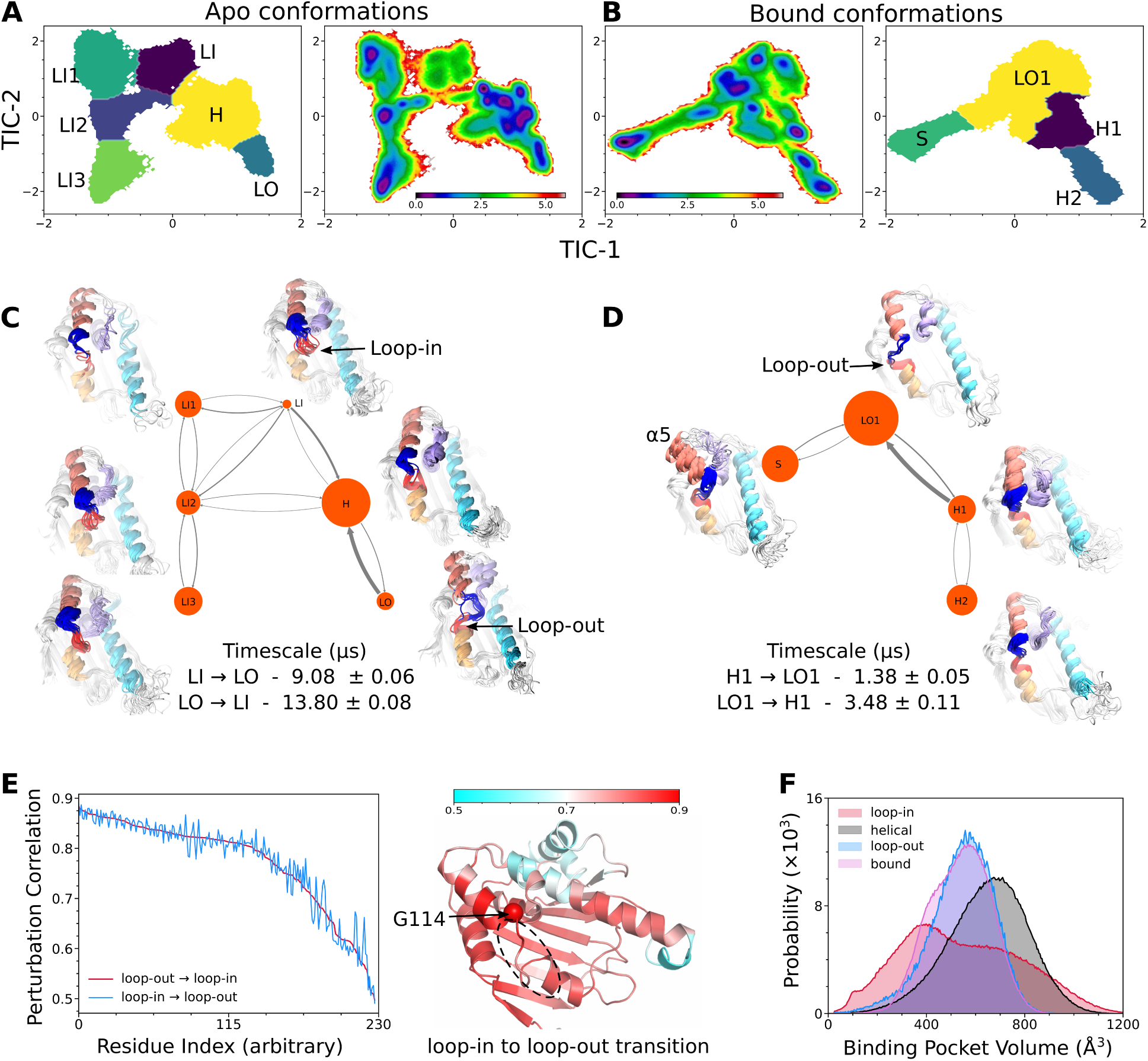
Conformational shift in N-Hsp90 by inhibitor binding. (A,B) The MSM weighted free energy surface and state maps for apo and bound states of N-Hsp90. (C,D) MSM models for apo and bound states. (E) Perturbation correlation estimated via PRS analysis for different transitions. Sphere represent G114 with highest correlation. Arbitrary residue index cor-repond to loop-out*→*loop-in decreasing order. (F) Binding pocket volumes of N-Hsp90 in different states (bootstrapped distribution of means).

To better understand the conformational landscape of the L4 region, it can be subdivided into two segments: residues 106-110 (red) and residues 111-115 (blue). In crystal structures, residues 106-110 face inward towards the binding pocket in the *loop-in* conformation and outward in the *loop-out* conformation. Representative snapshots from the six macrostates of the apo simulations (figure 3C) reveal the presence of both *loop-in* (LI, LI1, LI2, LI3; differing at *α*6-7 helices) and *loop-out* (LO) conformations. The remaining macrostates correspond to a helical intermediate orientation (H) of L4. These findings align with crystallographic observations, as the apo form of N-Hsp90 has been determined in various versions of *loop-out* and *loop-in* conformations. While the H state is not exactly helical, similar structures have been observed in crystal structures like 2XHR^46^ or 5J6N.^18^ The loop-in and helical states dominate in the apo form of N-Hsp90 compared to the loop-out conformation. These conformations are spontaneously interconvertible on microsecond timescales (figure S4).

To further validate these observations, we assessed whether the *loop-out* to *loop-in* and vice versa conformational transitions in the apo state could occur spontaneously. Perturbation response scanning (PRS) analysis^39^ was employed to investigate these transitions. In PRS, starting from N-Hsp90 in one conformation (e.g., *loop-out*), small random perturbations of force and direction were applied to each residue to determine whether the perturbations could drive N-Hsp90 into the *loop-in* conformation. If perturbations applied to most residues lead to the *loop-out* to *loop-in* transition, it suggests that the transition is spontaneous, and vice versa. PRS was performed for both *loop-in* to *loop-out* and *loop-out* to *loop-in* transitions, using the crystallographic poses of the *loop-out* (1YES) and *loop-in* (1YER) conformations of the apo form as reference end states. The perturbation correlation analysis indicated that all residues near the binding pocket exhibited similar and high correlations (*>*0.7). Thus, we concluded that the *loop-in* and *loop-out* conformations are spontaneously interconvertible, consistent with MSM timescales. The presence of all possible conformations of the L4 region in the apo state within short, achievable timescales suggests that GDM binding may proceed via a conformational selection mechanism, selectively binding and stabilizing the minor *loop-out* conformation, as observed in the GDM-bound crystallographic pose.

Next, we analyzed the conformational space of the substrate-bound state by constructing an MSM using the corresponding simulation ensemble, with the expectation of finding a predominant *loop-out* conformation. The four state converged macrostates derived from the MSM of the substrate-bound pose indicated the dominance of the *loop-out* conformation, with no *loop-in* conformation present. However, other helical-like states (H1, H2) were also observed (figure 3D). Importantly, the *loop-out* and helical conformations in the bound state were distinct from their counterparts in the apo state MSM. The representative snapshots in figures 3C and 3D correspond to the deepest free energy minima and do not represent the entire macrostate. A direct comparison of the free energy surface of the bound conformations with that of the apo conformations (figures 3A, B, S4B) reveals significant shifts in the positions of all minima, except for one helical macrostate. Therefore, the *loop-out* state in the bound form is labeled as LO1, and similarly, the helical states are labeled H1 and H2 (differing at *α*6-7). An additional state, S, appeared, which is entirely different from the apo states, characterized by a significant rotation of *α*5 away from *α*6-7. To our knowledge, while S has parallels in reported crystallographic poses (e.g., 1YES^24^ vs. 5J2V^18^), it does not exactly match any of the hundreds of crystallographic poses determined to date. These states are interconvertible on microsecond timescales but cannot be directly compared with their apo counterparts due to the differences observed. These observations suggest that while GDM binding shifts the equilibrium of the substrate-bound conformational space towards the *loop-out* state, it also generates new conformations and alternative versions of the *loop-out* and helical states, thereby supporting the induced-fit mechanism and challenging the conformational selection hypothesis.

Finally, we examined the widely held belief that the *loop-in* and *loop-out* conformations differ in their binding pocket volumes,^24^ with GDM binding to the *loop-out* conformation presumably having a larger binding pocket. However, the calculated binding pocket volumes across all conformational states in the apo MSM did not confirm this trend, as the *loop-out* state did not correspond to a larger pocket volume (figure 3F). Moreover, the pocket volume in the bound simulations also did not correlate with larger volumes. Overall, these results depict a complex picture of GDM-induced conformational changes in N-Hsp90, suggesting that both conformational selection and induced fit mechanisms may be at play.

### A Machine Learning Classifier Quantifies the Extent of Conformational Selection and Induced Fit in the Substrate Recognition Process

The preceding section indicates that GDM binding alters the conformational space of N-Hsp90 in a complex manner, suggesting potential involvement of both conformational selection and induced-fit mechanisms. This dual mechanism is not uncommon in biophysics; various systems, including proteins^47,48^ and RNA,^49^ have been shown to exhibit diverse combinations of conformational selection and induced-fit mechanisms. For GDM binding to Hsp90, we propose a similar scenario, where GDM binding initially follows a conformational selection pathway by selectively binding to the *loop-out* state of apo N-Hsp90. This binding is subsequently followed by conformational changes induced in the *loop-out* state, resulting in a different loop-out conformation (figure 4A). This phenomenon is referred to as “residual induced fit” in the literature.^50^ To explore this, we developed a machine learning classifier, specifically a Random Forest (RF) model, to exclusively detect the conformational changes associated with GDM binding through conformational selection and induced-fit pathways.

**Figure 4:**
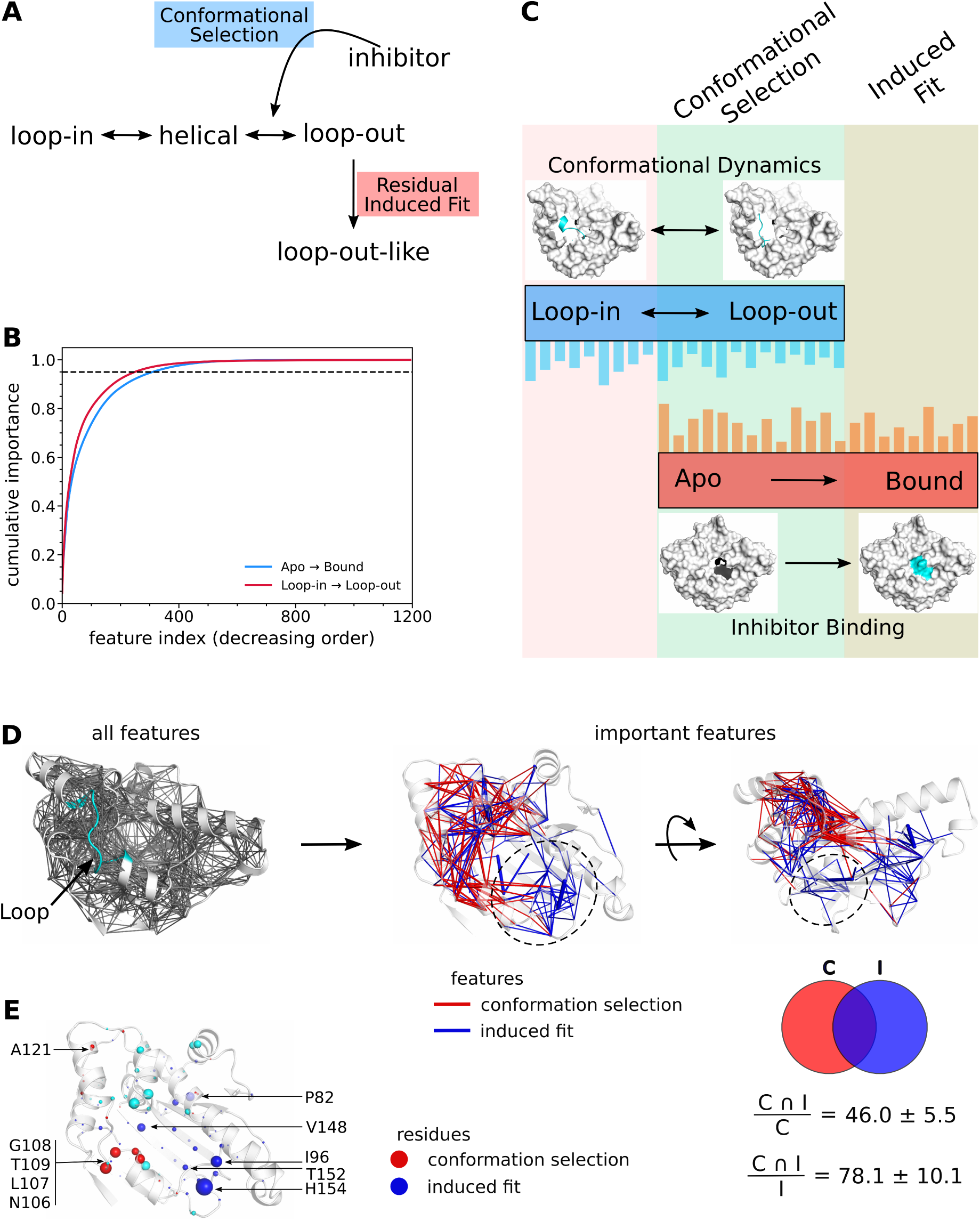
Conformation selection vs induced fit in N-Hsp90. (A) Residual induced fit scheme for Hsp90. (B) Mean cumulative feature importance scores obtained by trained RF classifiers. (C) Schematic for segregation of conformational selection and induced fit. (D) All and selected features based on feature importance scores and colored according to conformational selection (red) or induced fit(blue). (E) Important residues involved in conformational selection and induced fit mechanism.

In addition to learning labels, RF models can weigh input features based on their ability to classify these labels. These weights, known as ‘feature importance scores,’ have been used to identify features involved in protein allostery^51^ and residues differentiating MSM states.^52^ Similarly, we employed RF-based feature importance scores to detect significant conformational changes in N-Hsp90. To define the global conformation of N-Hsp90, native contacts (minimum distances in simulations) between residue pairs within 10 *Å* in the crystal structure were used. These contacts capture conformational changes across the entire protein, not just in the loop-4 region (as in MSM). Thus, each simulated conformation was represented by numerical values, which served as input features for RF training, with each feature representing a conformational change between two residues in terms of minimum distance.

In the first part of our analysis, input features from apo simulation trajectories were used, and each conformation was labeled with MSM-derived states: *loop-out*, *loop-in*, and helical, as shown in figure 3C. The RF models were successfully trained, achieving an accuracy of 0.99 *±* 0.00 (10e-7) and an F1 score of 0.98 *±* 0.00 (10e-4) (figure S5). Feature importance scores revealed that out of 1194 features, only 245*±*2 features corresponded to 95% of the conformational dynamics defining the MSM states (labels). Replicating the RF training on 50 cross-validated samples indicated that 206 of these 245*±*2 features consistently emerged as important across all samples (figure 4B). These 206 features represent the set of conformational changes that define the intrinsic dynamics of the apo states, referred to as ‘conf1,’ with their associated feature importance scores reflecting their relative contributions.

In the second part, another RF classifier was trained on input features from both apo and bound simulations. The input features were labeled as apo or bound (0 and 1), based on the simulations. The training was successful, with an accuracy of 0.99*±*0.00 and an F1 score of 0.99*±*0.00 (figure S5). According to the feature importance scores, 304*±*3 features out of 1194 accounted for 95% of the conformational difference between apo and bound states. Of these, 256 features were consistent across all 50 cross-validated replicates (figure 4B). These features represent the set of conformational changes that define the comparative dynamics between apo and bound states, i.e., the GDM binding process, referred to as ‘conf2.’

We hypothesized that the conformational changes associated with GDM binding (‘conf2’) that are also part of the intrinsic apo dynamics (‘conf1’) represent the features responsible for conformational selection, as these changes can spontaneously occur within the apo state (figure 4C). Conversely, the unique conformational changes in ‘conf2’ that are not observed in the apo state (‘conf1’) likely represent the features responsible for the induced-fit mechanism. The remaining conformational changes in ‘conf1’ that are not part of ‘conf2’ would not have any role in the GDM binding process. This allowed us to spatially locate and quantify the different conformational changes. Superimposing the conformational changes of ‘conf1’ and ‘conf2’ on the N-Hsp90 structure revealed an almost segregated presence of conformational selection (common in ‘conf1’ and ‘conf2’, shown in red) and the induced-fit mechanism (specific to ‘conf2’, shown in blue) (figure 4D). Interestingly, we found that the features responsible for conformational selection are specifically located in the L4 region, *α*5, *α*6, and *α*7, while the features responsible for induced fit are mostly located in the L3 and buried binding pocket regions (figure 4D). Summing the feature importance scores, 46.0*±*5.5% of the conformational changes in ‘conf1’ are common with ‘conf2,’ representing conformational selection, which accounts for 78.1*±*10.1% of ‘conf2.’ This suggests that the remaining 28% of ‘conf2’ features correspond to the induced-fit mechanism. As a further check, we repeated the analysis using input features only from loop-4 (as in MSM) and observed similar quantifications: 42.9*±*6.8% of ‘conf1’ represents 67.9*±*12.3% of ‘conf2’ conformational changes.

To make this analysis more meaningful at the residue level, we dissected the conformational changes further. Each line in figure 4D represents a feature, i.e., the minimum distance between two residues. The feature importance score of each feature was equally divided between its two residues, and these scores were summed to represent each residue’s relative importance. We observed three types of residues: (a) those associated only with the conformational selection route (red), (b) those associated only with induced fit (blue), and (c) those associated with both (cyan) (figure 4E). The residues predicted to be involved in conformational selection are exclusively located in the L4 region (N106, L107, G108, T109), which is also the most extensively investigated region of N-Hsp90. In particular, L107 has recently been identified as the most important residue for its entropic contribution.^18^ This is consistent with previous reports that the L4 region exhibits conformational heterogeneity within the apo state, with inhibitors specifically binding to some of these conformations.^18^ The residues predicted to be involved in the induced-fit mechanism are primarily located in the L3 region and within the binding pocket. Coincidentally, this L3 region is where a binding intermediate was observed (figure 1), validating the existence of an intermediate state. This observation aligns with previous reports suggesting the existence of an intermediate state and an induced-fit pathway,^16^ based on the inference that the induced-fit mechanism proceeds through intermediate steps. Notably, the simulations containing the intermediate state were not used in the above analysis, yet induced fit in the L3 region was independently observed. The cyan residues in *α*6-7 may either be involved in both conformational selection and induced-fit mechanisms or reflect limitations in the methodology employed in this study.

## Conclusion

In this study, we present a detailed analysis of the inhibitor binding mechanism to human N-Hsp90 and report the identification of a putative intermediate state. This intermediate state was observed through unbiased binding simulations, which were conducted without any prior bias towards the binding site. These simulations are well-regarded for their ability to recover binding modes accurately.^23,43,44^ Our findings are further supported by the convergence of Markov State Models (MSMs) to two or three states, rather than a larger number, reinforcing the notion that inhibitor binding to N-Hsp90 occurs through a two-step process.

The generalizability of our findings is based on the assumption that other known inhibitors will follow a similar binding mechanism as observed for GDM. Given the extensive variety of GDM derivatives and the large number of potential inhibitors, it is impractical to study each individually. Therefore, we focused on the parent molecule, GDM, which serves as a representative for many inhibitors currently under design. Since these inhibitors target the same ATP binding site, we anticipate that their binding mechanisms will be analogous to those reported here for GDM.

Our results reveal that inhibitor binding induces significant changes in the conformational space of Hsp90, which can be attributed to the simultaneous presence of both conformational selection and induced fit mechanisms. This study provides a quantitative estimate of the extents to which these mechanisms contribute to inhibitor binding. The coexistence of these two binding regimes is not uncommon, ^47–50^ yet it remains challenging to estimate their relative contributions accurately.^53^ To the best of our knowledge, no existing computational methods are available to achieve this level of detail.

Furthermore, our work includes a detailed residue-level analysis of conformational selection and induced fit mechanisms, which is crucial for advancing inhibitor design. Our findings are consistent with previous studies that have identified the loop-4 region as being involved in the conformational selection process through crystallography^24^ and biochemical assays.^18^ This study supports and expands upon these previous findings by demonstrating the mutual involvement of both conformational selection and induced fit in the inhibitor binding process.

## Supplementary Information

figures S1-S5.

## Data and code availability

Most of the data and all codes are available at https://github.com/msahilgit/hsp90

## Acknowledgement

We acknowledge support of the Department of Atomic Energy, Government of India, under Project Identification No. RTI 4007. JM acknowledges Core Research grants provided by the Department of Science and Technology (DST) of India (CRG/2023/001426). We thank Dr. Sneha Menon and Subinoy Adhikari for discussions and their useful suggestions. This work was supported by shared computing resources obtained from TIFR Hyderabad, India.

## References

(1) Zhang, J.; Li, H.; Liu, Y.; Zhao, K.; Wei, S.; Sugarman, E. T.; Liu, L.; Zhang, G. Targeting HSP90 as a novel therapy for cancer: mechanistic insights and translational relevance. Cells 2022, 11, 2778.

(2) Hartl, F. U.; Bracher, A.; Hayer-Hartl, M. Molecular chaperones in protein folding and proteostasis. Nature 2011, 475, 324–332.

(3) Karagöz, G. E.; Rüdiger, S. G. Hsp90 interaction with clients. Trends in biochemical sciences 2015, 40, 117–125.

(4) Schopf, F. H.; Biebl, M. M.; Buchner, J. The HSP90 chaperone machinery. Nature reviews Molecular cell biology 2017, 18, 345–360.

(5) Biebl, M. M.; Buchner, J. Structure, function, and regulation of the Hsp90 machinery. Cold Spring Harbor perspectives in biology 2019, 11, a034017.

(6) Castelli, M.; Magni, A.; Bonollo, G.; Pavoni, S.; Frigerio, F.; Oliveira, A. S. F.; Cinquini, F.; Serapian, S. A.; Colombo, G. Molecular mechanisms of chaperone-directed protein folding: Insights from atomistic simulations. Protein Science 2024, 33, e4880.

(7) Jaeger, A. M.; Whitesell, L. HSP90: enabler of cancer adaptation. Annual Review of Cancer Biology 2019, 3, 275–297.

(8) Whitesell, L.; Lindquist, S. L. HSP90 and the chaperoning of cancer. Nature Reviews Cancer 2005, 5, 761–772.

(9) Whitesell, L.; Shifrin, S. D.; Schwab, G.; Neckers, L. Benzoquinonoid ansamycins possess selective tumoricidal activity unrelated to src kinase inhibition. Cancer research 1992, 52, 1721–1728.

(10) Ardestani, M.; Khorsandi, Z.; Keshavarzipour, F.; Iravani, S.; Sadeghi-Aliabadi, H.; Varma, R. S. Heterocyclic compounds as Hsp90 inhibitors: a perspective on anticancer applications. Pharmaceutics 2022, 14, 2220.

(11) Li, Z.-N.; Luo, Y. HSP90 inhibitors and cancer: Prospects for use in targeted therapies. Oncology Reports 2023, 49, 1–13.

(12) Khandelwal, A.; Crowley, V. M.; Blagg, B. S. Natural product inspired N-terminal Hsp90 inhibitors: from bench to bedside? Medicinal research reviews 2016, 36, 92–118.

(13) Skrzypczak, N.; Buczkowski, A.; Bohusz, W.; Nowak, E.; Tokarska, K.; Lésniewska, A.; Alzebari, A. M.; Ruszkowski, P.; Gdaniec, M.; Bartl, F., et al. Modifications of gel-danamycin via CuAAC altering affinity to chaperone protein Hsp90 and cytotoxicity. European Journal of Medicinal Chemistry 2023, 256, 115450.

(14) Li, Z.; Jia, L.; Tang, H.; Shen, Y.; Shen, C. LZY3016, a novel geldanamycin derivative, inhibits tumor growth in an MDA-MB-231 xenograft model. RSC advances 2023, *13*, 13586–13591.

(15) Wang, X.; Zhang, Y.; Ponomareva, L. V.; Qiu, Q.; Woodcock, R.; Elshahawi, S. I.; Chen, X.; Zhou, Z.; Hatcher, B. E.; Hower, J. C., et al. Mccrearamycins A– D, Geldanamycin-Derived Cyclopentenone Macrolactams from an Eastern Kentucky Abandoned Coal Mine Microbe. Angewandte Chemie 2017, 129, 3040–3044.

(16) Gooljarsingh, L. T.; Fernandes, C.; Yan, K.; Zhang, H.; Grooms, M.; Johanson, K.; Sinnamon, R. H.; Kirkpatrick, R. B.; Kerrigan, J.; Lewis, T., et al. A biochemical rationale for the anticancer effects of Hsp90 inhibitors: slow, tight binding inhibition by geldanamycin and its analogues. Proceedings of the National Academy of Sciences 2006, 103, 7625–7630.

(17) Onuoha, S.; Mukund, S.; Coulstock, E.; Sengerova, B.; Shaw, J.; McLaughlin, S.; Jackson, S. Mechanistic studies on Hsp90 inhibition by ansamycin derivatives. Journal of molecular biology 2007, 372, 287–297.

(18) Amaral, M.; Kokh, D.; Bomke, J.; Wegener, A.; Buchstaller, H.; Eggenweiler, H.; Matias, P.; Sirrenberg, C.; Wade, R.; Frech, M. Protein conformational flexibility modulates kinetics and thermodynamics of drug binding. Nature communications 2017, 8, 2276.

(19) Zhang, H.; Zhou, C.; Chen, W.; Xu, Y.; Shi, Y.; Wen, Y.; Zhang, N. A dynamic view of ATP-coupled functioning cycle of Hsp90 N-terminal domain. Scientific reports 2015, 5, 9542.

(20) Hellemann, E.; Durrant, J. D. Worth the Weight: Sub-Pocket EXplorer (SubPEx), a Weighted Ensemble Method to Enhance Binding-Pocket Conformational Sampling. Journal of Chemical Theory and Computation 2023, 19, 5677–5689.

(21) Sohmen, B.; Beck, C.; Frank, V.; Seydel, T.; Hoffmann, I.; Hermann, B.; Nüesch, M.; Grimaldo, M.; Schreiber, F.; Wolf, S., et al. The Onset of Molecule-Spanning Dynamics in Heat Shock Protein Hsp90. Advanced Science 2023, 10, 2304262.

(22) Colombo, G.; Morra, G.; Meli, M.; Verkhivker, G. Understanding ligand-based modulation of the Hsp90 molecular chaperone dynamics at atomic resolution. Proceedings of the National Academy of Sciences 2008, 105, 7976–7981.

(23) Sahil, M.; Singh, T.; Ghosh, S.; Mondal, J. 3site Multisubstrate-Bound State of Cytochrome P450cam. Journal of the American Chemical Society 2023, 145, 23488–23502.

(24) Stebbins, C. E.; Russo, A. A.; Schneider, C.; Rosen, N.; Hartl, F. U.; Pavletich, N. P. Crystal structure of an Hsp90–geldanamycin complex: targeting of a protein chaperone by an antitumor agent. Cell 1997, 89, 239–250.

(25) Moroni, E.; Morra, G.; Colombo, G. Molecular dynamics simulations of Hsp90 with an eye to inhibitor design. Pharmaceuticals 2012, 5, 944–962.

(26) Nazar, A.; Abbas, G.; Azam, S. S. Deciphering the inhibition mechanism of under trial Hsp90 inhibitors and their analogues: a comparative molecular dynamics simulation. Journal of Chemical Information and Modeling 2020, 60, 3812–3830.

(27) Huang, J.; MacKerell Jr, A. D. CHARMM36 all-atom additive protein force field: Validation based on comparison to NMR data. J. Comput. Chem. 2013, 34, 2135– 2145.

(28) Huang, L.; Roux, B. Automated force field parameterization for nonpolarizable and polarizable atomic models based on ab initio target data. J. Comput. Theory Chem. 2013, 9, 3543–3556.

(29) Bussi, G.; Donadio, D.; Parrinello, M. Canonical sampling through velocity rescaling. The Journal of chemical physics 2007, 126.

(30) Parrinello, M.; Rahman, A. Polymorphic transitions in single crystals: A new molecular dynamics method. Journal of Applied physics 1981, 52, 7182–7190.

(31) Van Der Spoel, D.; Lindahl, E.; Hess, B.; Groenhof, G.; Mark, A. E.; Berendsen, H. J. GROMACS: fast, flexible, and free. J. Comput. Chem. 2005, 26, 1701–1718.

(32) Darden, T.; York, D.; Pedersen, L. Particle mesh Ewald: An NlogN method for Ewald sums in large systems. J. Chem. Phys. 1993, 98, 10089–10092.

(33) Pail, S.; Hess, B. A flexible algorithm for calculating pair interactions on SIMD architectures. Comput. Phys. Commun. 2013, 184, 2641–2650.

(34) Hess, B.; Bekker, H.; Berendsen, H. J.; Fraaije, J. G. LINCS: a linear constraint solver for molecular simulations. J. Comput. Chem. 1997, 18, 1463–1472.

(35) Miyamoto, S.; Kollman, P. A. Settle: An analytical version of the SHAKE and RATTLE algorithm for rigid water models. J. Comput. Chem. 1992, 13, 952–962.

(36) Husic, B. E.; Pande, V. S. Markov state models: From an art to a science. Journal of the American Chemical Society 2018, 140, 2386–2396.

(37) Scherer, M. K.; Trendelkamp-Schroer, B.; Paul, F.; Pérez-Hernández, G.; Hoffmann, M.; Plattner, N.; Wehmeyer, C.; Prinz, J.-H.; Nóe, F. PyEMMA 2: A software package for estimation, validation, and analysis of Markov models. Journal of chemical theory and computation 2015, 11, 5525–5542.

(38) Harrigan, M. P.; Sultan, M. M.; Herández, C. X.; Husic, B. E.; Eastman, P.; Schwantes, C. R.; Beauchamp, K. A.; McGibbon, R. T.; Pande, V. S. MSMBuilder: statistical models for biomolecular dynamics. Biophysical journal 2017, 112, 10–15.

(39) Penkler, D.; Sensoy, O.; Atilgan, C.; Tastan Bishop, O. Perturbation–response scanning reveals key residues for allosteric control in Hsp70. Journal of Chemical Information and Modeling 2017, 57, 1359–1374.

(40) Brown, D. K.; Penkler, D. L.; Sheik Amamuddy, O.; Ross, C.; Atilgan, A. R.; Atilgan, C.; Tastan Bishop, Ö. MD-TASK: a software suite for analyzing molecular dy-namics trajectories. Bioinformatics 2017, 33, 2768–2771.

(41) Schmidtke, P.; Bidon-Chanal, A.; Luque, F. J.; Barril, X. MDpocket: open-source cavity detection and characterization on molecular dynamics trajectories. Bioinformatics 2011, 27, 3276–3285.

(42) Tanemura, M.; Ogawa, T.; Ogita, N. A new algorithm for three-dimensional Voronoi tessellation. J. Comput. Phys. 1983, 51, 191–207.

(43) Greisman, J. B.; Willmore, L.; Yeh, C. Y.; Giordanetto, F.; Shahamadtar, S.; Nisonoff, H.; Maragakis, P.; Shaw, D. E. Discovery and validation of the binding poses of allosteric fragment hits to protein tyrosine phosphatase 1b: From molecular dynamics simulations to X-ray crystallography. Journal of Chemical Information and Modeling 2023, 63, 2644–2650.

(44) Sahil, M.; Singh, J.; Sahu, S.; Pal, S. K.; Yadav, A.; Anand, R.; Mondal, J. Identifying Selectivity Filters in Protein Biosensor for Ligand Screening. JACS Au 2023, 3, 2800– 2812.

(45) Motlagh, H. N.; Wrabl, J. O.; Li, J.; Hilser, V. J. The ensemble nature of allostery. Nature 2014, 508, 331–339.

(46) Murray, C. W.; Carr, M. G.; Callaghan, O.; Chessari, G.; Congreve, M.; Cowan, S.; Coyle, J. E.; Downham, R.; Figueroa, E.; Frederickson, M., et al. Fragment-based drug discovery applied to Hsp90. Discovery of two lead series with high ligand efficiency. Journal of medicinal chemistry 2010, 53, 5942–5955.

(47) Wang, Q.; Zhang, P.; Hoffman, L.; Tripathi, S.; Homouz, D.; Liu, Y.; Waxham, M. N.; Cheung, M. S. Protein recognition and selection through conformational and mutually induced fit. Proceedings of the National Academy of Sciences 2013, 110, 20545–20550.

(48) Ewers, D.; Becher, T.; Machtens, J.-P.; Weyand, I.; Fahlke, C. Induced fit substrate binding to an archeal glutamate transporter homologue. Proceedings of the National Academy of Sciences 2013, 110, 12486–12491.

(49) Suddala, K. C.; Wang, J.; Hou, Q.; Walter, N. G. Mg2+ shifts ligand-mediated folding of a riboswitch from induced-fit to conformational selection. Journal of the American Chemical Society 2015, 137, 14075–14083.

(50) Wlodarski, T.; Zagrovic, B. Conformational selection and induced fit mechanism underlie specificity in noncovalent interactions with ubiquitin. Proceedings of the National Academy of Sciences 2009, 106, 19346–19351.

(51) Ahalawat, N.; Sahil, M.; Mondal, J. Resolving Protein Conformational Plasticity and Substrate Binding via Machine Learning. Journal of Chemical Theory and Computation 2023, 19, 2644–2657.

(52) Westerlund, A. M.; Sridhar, A.; Dahl, L.; Andersson, A.; Bodnar, A.-Y.; Delemotte, L. Markov state modelling reveals heterogeneous drug-inhibition mechanism of Calmodulin. PLoS Computational Biology 2022, 18, e1010583.

(53) Paul, F.; Weikl, T. R. How to distinguish conformational selection and induced fit based on chemical relaxation rates. PLoS computational biology 2016, 12, e1005067.

