## Supplemental figures and table for "Dynamic Insights into Hsp90 Inhibitor Binding: Uncovering Intermediate States and Conformational Plasticity"

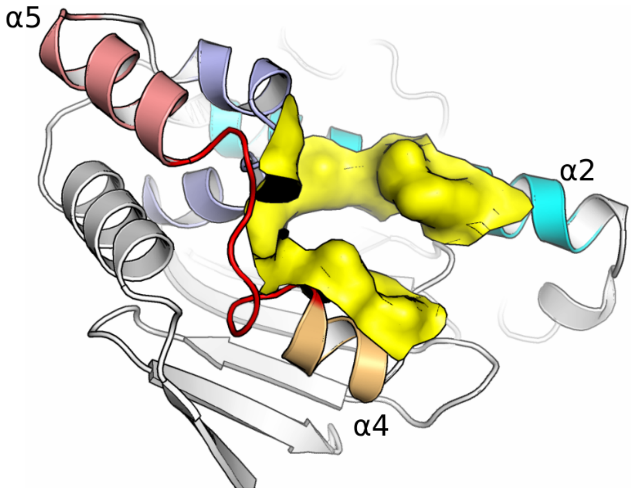


**Figure S1:** The binding pocket residues (yellow surface) used to measure ligand distance on-the-fly to detect binding events.

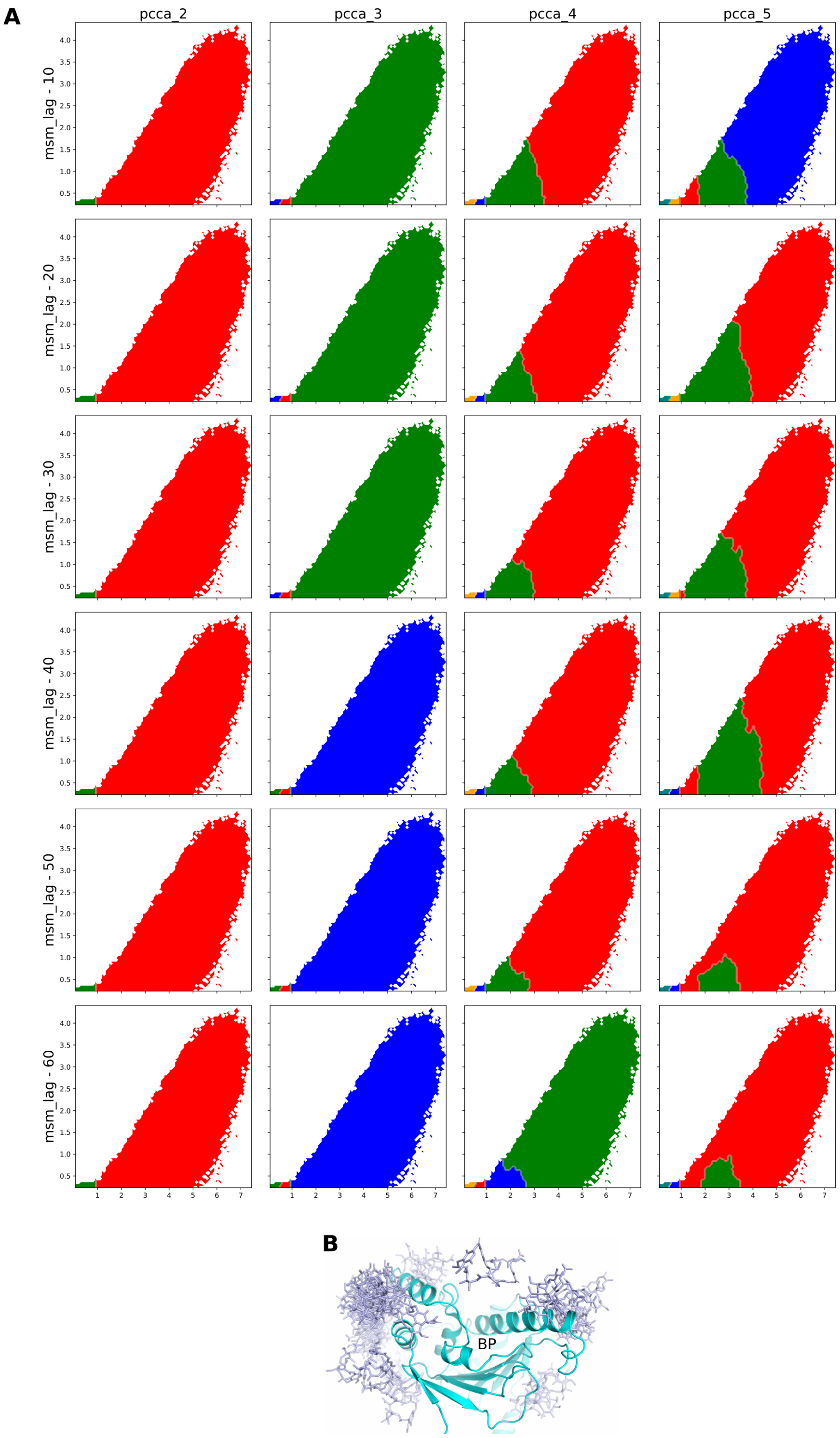


**Figure S2:** (A) MSM state maps for different number of pcca states at various lag times. (B) The fourth state observed in four state MSM with respect to binding pocket. The other states were unbound, intermediate and bound states similar to three state MSM.

| state | population |
|---|---|
| U | 0.40 ± 0.02 |
| I | 0.08 ± 0.002 |
| B | 0.52 ± 0.02 |

| transition | mfpt (μs) |
|---|---|
| U → I | 26.7 ± 3.2 |
| I → B | 1.5 ± 0.06 |
| U → B | 35.3 ± 2.7 |

**Figure S3:** Equilibrium state populations and mean first passage transition times for different binding states as estimated by MSM.

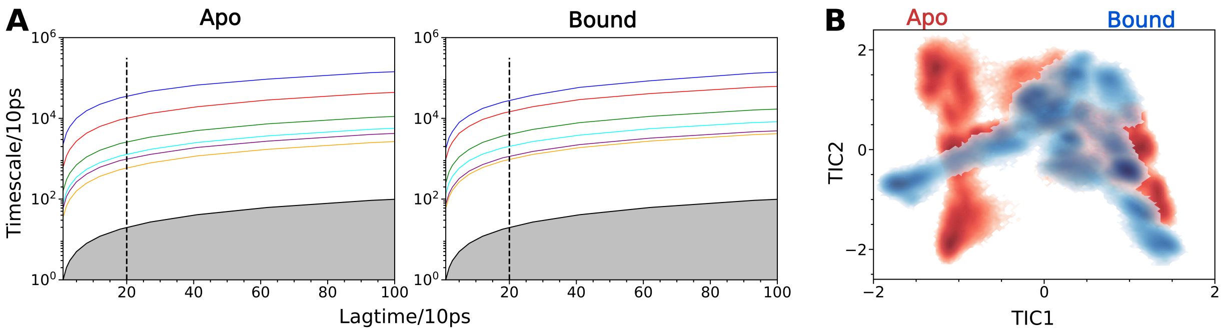


**C**

| | C | C2 | O | C1 | C3 | I |
|---|---|---|---|---|---|---|
| C | 0 | 7.55±0.21 | 9.08±0.06 | 12.7±1.10 | 20.88±3.09 | 4.58±0.03 |
| C2 | 13.61±0.10 | 0 | 15.04±0.12 | 7.20±1.74 | 10.47±2.77 | 10.6±0.10 |
| O | 13.80±0.08 | 13.38±0.17 | 0 | 20.18±1.32 | 27.12±3.04 | 0.19±0.00 |
| C1 | 12.67±0.15 | 3.26±0.55 | 15.61±0.09 | 0 | 14.57±3.46 | 11.16±0.07 |
| C3 | 16.42±0.65 | 2.06±0.59 | 17.93±0.50 | 9.51±2.28 | 0 | 13.49±0.52 |
| I | 13.29±0.08 | 12.88±0.17 | 3.54±0.03 | 19.68±1.32 | 26.62±3.04 | 0 |

| | I1 | I2 | S | O1 |
|---|---|---|---|---|
| I1 | 0 | 21.2±0.18 | 13.43±1.62 | 1.38±0.05 |
| I2 | 4.25±0.00 | 0 | 17.46±1.62 | 5.27±0.09 |
| S | 9.36±0.50 | 32.1±0.60 | 0 | 3.29±0.78 |
| O1 | 3.48±0.11 | 26.24±0.28 | 10.53±1.50 | 0 |

**D**

| | C | C2 | O | C1 | C3 | I |
|---|---|---|---|---|---|---|
| | 1.51±0.01 | 11.92±0.17 | 6.4±0.02 | 14.53±0.17 | 16.3±0.32 | 49.34±0.40 |

| | I1 | I2 | S | O1 |
|---|---|---|---|---|
| | 12.91±0.39 | 15.81±0.12 | 23.9±1.47 | 47.38±1.57 |

**Figure S4:** (A) The ITS plots for MSM models of apo and bound states. (B) Overlay of free energy surfaces apo and bound MSM. (C,D) Mean first passage transition times and equilibrium populations of different states of apo and bound MSMs.

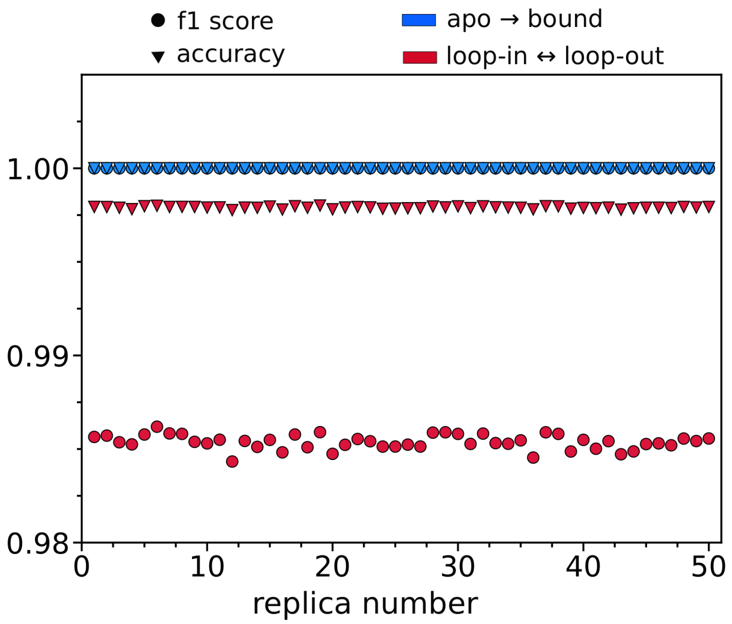


**Figure S5:** The accuracy and f1 scores for different replicas of trained RF on test data.
